# Accuracy of short tandem repeats genotyping tools in whole exome sequencing data

**DOI:** 10.1101/2020.02.03.933002

**Authors:** Andreas Halman, Alicia Oshlack

## Abstract

**Background:** Short tandem repeats are important source of genetic variation, they are highly mutable and repeat expansions are associated dozens of human disorders, such as Huntington’s disease and spinocerebellar ataxias. Technical advantages in sequencing technology have made it possible to analyse these repeats at large scale, however, accurate genotyping is still a challenging task. We compared four different short tandem repeats genotyping tools on whole exome sequencing data to determine their genotyping performance and limits which will aid other researchers to choose a suitable tool and parameters for analysis.

**Methods:** The analysis was performed on the Simons Simplex Collection dataset where we used a novel method of evaluation with accuracy determined by the rate of homozygous calls on the X chromosome of male samples. In total we analysed 433 samples and around a million genotypes for evaluating tools on whole exome sequencing data.

**Results:** We determined a relatively good performance of all tools when genotyping repeats of 3-6 bp in length which could be improved with coverage and quality score filtering. However, genotyping homopolymers was challenging for all tools and a high error rate was present across different thresholds of coverage and quality scores. Interestingly, dinucleotide repeats displayed a high error rate as well, which was found to be mainly caused by the AC/TG repeats. Overall, LobSTR was able to make the most calls and was also the fastest tool while RepeatSeq and HipSTR exhibited the lowest heterozygous error rate at low coverage.

**Conclusions:** All tools have different strengths and weaknesses and the choice may depend on the type of analysis. In this analysis we demonstrated the effect of using different filtering parameters and offered recommendations based on the trade-off between the best accuracy of genotyping and the highest number of calls.

## Introduction

### Overview of short tandem repeats and methods of analysis

Short tandem repeats (STRs), also known as microsatellites, consist of repeated units of 1 to 6 base pairs (bp) in length and cover about 3% of the human genome (Gymrek, 2017). STRs are highly mutable and often vary in their number of repeat units across the population. They can be found in various regions of the genome, including in or near protein coding regions and introns (Hannan, 2018). Expanded variants contribute to several dozen human disorders, including Huntington’s disease, fragile X syndrome, spinocerebellar ataxias and other diseases. In addition, variation in STR length has been shown to associate with quantitative traits such as gene expression (Gymrek, 2017). The standard method to genotype the length of STRs is to perform polymerase chain reaction (PCR) amplification on the region of interest and gel electrophoresis (Tang, Haixu et al., 2017). Sanger sequencing has high accuracy, but low throughput, limiting analysis to a few genes at a time (Caspar et al., 2018).

Recent technology advances in high-throughput sequencing (HTS) has revolutionised genomics field and brought us the opportunity to detect sequence variants at a scale that was impossible before (Caspar et al., 2018). HTS of the whole genome provides the potential to profile over a million STRs in the human genome. Recent advances in bioinformatics have brought us tools to analyse STRs from sequencing data (HTS), but genotyping still remains challenging for many reasons, including issues with extreme GC content, short read lengths that do not span over the entire repeat, and issues with alignment due to variation in STRs appearing as large insertions or deletions relative to the reference. In addition, using PCR amplification during library preparation will often cause stutter noise and produce artificial variability in the sequence (Gymrek, 2017; Caspar et al., 2018). Stutter noise is a result of *in vitro* slippage of DNA polymerase during PCR cycles that leads to erroneous reads of incorrect repeat length (Willems et al., 2014) which contributes to challenges in genotyping.

Illumina has developed a method for amplification-free (PCR−) library preparation (Kozarewa et al., 2009) which theoretically eliminates the STR stutter error during PCR amplification in sample preparation (PCR+) and therefore improves the accuracy of STR genotyping. The developers of STR-FM evaluated the new protocol by running their tool in both PCR− and PCR+ samples and found that PCR− protocol compared to PCR+ has up to 9-fold fewer errors (Fungtammasan et al., 2015). However, huge amounts of sequencing data have already been generated by using the PCR+ protocol where some data will not be resequenced due to time and/or cost (Fungtammasan et al., 2015). In addition, despite the advantages of whole genome sequencing (WGS), whole exome sequencing (WES) is still widely used in human genetics due to its lower cost and higher coverage and WES is a PCR+ process (Björn et al., 2018). Therefore, tools that can accurately genotype STRs not only from PCR− but also from PCR+ data are essential.

While there are a number of computational tools that have been developed to genotype STR alleles in HTS data there have been few independent comparisons of their performance. Evaluation of methods for genotyping STRs is difficult. The gold standard measurement of STRs is by capillary electrophoresis (Gymrek, 2016) but these methods have low throughput. Further evaluations have used Mendelian inheritance as a measure of accuracy (Gymrek et al., 2012; Highnam et al., 2013; Mousavi et al., 2019). Other studies have used simulated data for the evaluation of genotyping accuracy (Highnam et al., 2013; Fungtammasan et al., 2015). While simulation can generate many loci with known alleles it is difficult to simulate the true complexity of real data.

Here we propose to compare and evaluate STR genotyping methods on exome data using a different but complementary approach. We used the natural hemizygous state of the X chromosomes in males to look for incorrect calls revealed by a heterozygous call. With repeats on the X chromosome in males there is only one allele so we expect all calls to be homozygous. While this approach does not evaluate the accuracy of the allele length it has advantages in that (a) the data sets are large so we can test thousands of calls (b) the data comes from real patients with all the noise and biases found in real data.

In our study, we compared LobSTR (Gymrek et al., 2012), RepeatSeq (Highnam et al., 2013), HipSTR (Willems et al., 2017) and a recently published tool GangSTR (Mousavi et al., 2019). In addition, we included a common variant calling tool GATK HaplotypeCaller (McKenna et al., 2010) as a comparison of genotyping accuracy.

There are a number of tools which have been developed for STR analysis and which were excluded from this analysis. For example, popSTR (Kristmundsdóttir et al., 2017) is a population based STR genotyper and optimised for whole genome sequencing (WGS) data. STRviper is another method for genotyping STRs that is able to pick up repeats longer than the read length, however, it has no built-in stutter model and it is not suitable for diploid dataset as it assumes only one allele (Cao et al., 2014). Galaxy environment has a STR analysis tool called STR-FM which we were unable to run (Fungtammasan et al., 2015). Dante (Budiš et al., 2019) and STRScan (Tang, Haixu et al., 2017) are designed for targeted searches and requires a user-defined list of STR loci.

Tools such as Expansion Hunter (Dolzhenko et al., 2017; Dolzhenko et al., 2019), TREDPARSE (Tang, Haibao et al., 2017), STRetch (Dashnow et al., 2018) and exSTRa (Tankard et al., 2018) were excluded from our analysis as well because they are classified as tools which specifically look for expansions that might be disease causing and are often longer than the physical read length or expansion relative to a control set.

Our analysis focuses on comparing the performance of STR genotyping tools on the X chromosome of more than 400 males. Using this data set we investigate the overall ability of tools to call genotypes, the accuracy as a function of coverage and repeat unit and also investigate quality scores of the tools. We find most tools are able to call a majority of homozygous alleles and different tools have different advantages in terms of repeat unit and coverage.

### Computational tools to genotype STRs from HTS data

First, we will give a short overview of STR genotyping tools included in our analysis and their reported accuracy. The tools evaluated in our analysis are summarised in Table 1. All of the tools require a set of defined STR loci. Tandem Repeats Finder (TRF) is a tool that can be used to detect STRs that have two or more copies of the same repeat unit in a row in the reference genome (Benson, 1999). Also, it can detect repeats for which the repeat unit size is up to several hundreds of bp long. Running TRF generates a report which includes all the loci detected in the genome with genomic start and end location of the STR, repeat unit and its size, number of copies aligned with the consensus pattern and other relevant information. For this study we limited the loci defined by TRF to repeat units up to 6 bp (see Methods).

**Table 1.** Feature comparison of STR specific genotyping tools used in our analysis.

|  | **RepeatSeq** | **LobSTR** | **HipSTR** | **GangSTR** |
| --- | --- | --- | --- | --- |
| **Latest version of the tool used in this study** | 0.8.2 (2014) | 4.0.6 (2016) | 0.6.2 (2018) | 2.4 (2019) |
| **Built-in stutter noise model** | ✓ | ✓ | ✓ | ✓ |
| **Ability to detect STRs that are longer than the read length** | ✗ | ✗ | ✗ | ✓ |
| **Input file types** | BAM | BAM<br>FASTA<br>FASTQ | BAM<br>CRAM | BAM<br>CRAM |
| **Sequencing read types** | Single- and<br>paired-end<br>reads | Single- and<br>paired-end<br>reads | Single- and<br>paired-end<br>reads | Paired-end<br>reads |

### LobSTR

LobSTR was one of the first successful STR genotyping tools for HTS data. It initially used its own inbuilt aligner but can also use data aligned with BWA-MEM (Li, 2013). LobSTR identifies reads that completely contain the STR and which also have flanking sequence with no repetitive sequence when aligned to a reference genome. As mentioned, PCR amplification during library preparation can create stutter noise at an STR locus, and LobSTR tackles this issue with an included stutter model that aims to detect and account for noise to improve genotyping accuracy. The stutter noise model can be custom generated from the data or use the standardised one supplied by the tool developers. As a result, LobSTR determines and reports the maximum likelihood estimates of the genotype in each locus. (Gymrek et al., 2012)

LobSTR was validated using concordance of biological replicates (blood and saliva samples) from the same subject to measure the precision of the tool. At 21x coverage the discordance rate for genotype was 3% and for allelotype 2%. While for lower, 5x coverage the discordance rate for genotype was 11% and for allelotype 5%. STR length differences were analysed in discordant calls that were heterozygous in both blood and saliva samples and found that at coverage 5x or higher, 90% of the errors were one repeat unit difference and 99% of errors were in 2 bp repeat unit size. (Gymrek et al., 2012) However, it is important to note that LobSTR validated 2-6 bp repeat unit size STRs and did not validate homopolymers (Gymrek et al., 2012), which are common in the genome and a known source of genetic variation (Highnam et al., 2013).

### RepeatSeq

RepeatSeq (Highnam et al., 2013) uses data aligned by external tool, such as BWA or Bowtie. It uses Bayesian model selection to determine the most probable genotype and requires all reads to fully contain STRs and at least two reads at a locus to make a call. The RepeatSeq noise model is based on genomes derived from over 100 inbred isolates of fly.

RepeatSeq’s accuracy was evaluated by analysing a trio WGS data to test consistency with Mendelian inheritance. The authors reported that on minimum coverage of two, 92.1% of repeat calls were consistent with the Mendelian inheritance, while with a minimum coverage of 9 it was 95.3% and on minimum coverage of 17 it was 98.0%. (Highnam et al., 2013)

### GangSTR

One of the major drawbacks of the first series of STR profiling methods was that they were limited to genotyping repeats within the read length in HTS data. GangSTR (Mousavi et al., 2019) is a more recent method that incorporates additional information besides repeat-enclosing reads to estimate the length of repeats. This includes available information such as fragment length, coverage and information about partially enclosing reads where only one end contains flanking sequence. More specifically, reads are divided into four classes: 1) enclosing read pairs that have at least one read which includes the whole STR and a flanking region in both ends; 2) spanning read pairs that have a mate pair where one read is aligned to one side of the STR and the second read of the pair on the other side; 3) flanking read pairs includes a read which partially extends into the STR region; 4) fully repetitive read pairs have one or two reads which are entirely made of STR (Mousavi et al., 2019). These four classes of reads are used to not only genotype repeats less than the read length but can also be used to genotype longer alleles such as repeat expansions.

The GangSTR method was evaluated by first simulating paired-end 150 bp reads (40x coverage) for 14 repeat expansions involved in STR disorders. Tool accuracy was measured by comparing true and observed alleles and also compared to TREDPARSE and ExpansionHunter. In this evaluation GangSTR showed lower root mean square error (RMSE) rate between true and observed allele lengths for all tested repeats. The authors demonstrated that GangSTR had an advantage over ExpansionHunter and TREADPARSE, especially in alleles which were close to the read length or longer. Also, GangSTR and ExpansionHunter improved significantly with higher coverage and longer read length.

GangSTR genotyping for disease causing alleles was also tested on validated 14 Huntington’s Disease and 25 Fragile X Syndrome real PCR−free WGS data and they reported of RMSE (7.9 and 29.3, respectively) that was lower than for TREDPARSE (8.3 and 34.8, respectively) and ExpansionHunter (10.1 and 27.3, respectively). In evaluations of genotyping a WGS trio, GangSTR was found to have similar performance to HipSTR for shorter alleles. (Mousavi et al., 2019)

### HipSTR

HipSTR (Willems et al., 2017) is a haplotype-based method for genotyping, haplotyping and phasing STRs. While other STR tools are made for finding true length of repeats independently along the genome, HipSTR takes into account the whole repeat structure on the allele which may also have missing data. HipSTR accuracy was tested by comparing calls from 118 PCR− WGS samples to capillary electrophoresis data, reporting about 98.8% consistency between the two datasets. (Willems et al., 2017)

### GATK HaplotypeCaller

GATK HaplotypeCaller (GATK-HC) (McKenna et al., 2010) can also be used for finding SNPs and indels in repeat regions, but it is not specifically made for STR analysis. It has been widely documented that indel calling is not as accurate as SNP calling and indel callers are not ideal for identifying STR mutation due to the lack of reporting repeat genotypes. Instead, indel callers report insertions or deletions of bases relative to the reference which may or may not be a multiple of the repeat unit as well as including SNP differences. Dedicated STR callers however use information about the repeat unit, composition and repeat length in order to make more accurate genotype calls. (Highnam et al., 2013)

## Results

In order to evaluate the accuracy and performance of STR genotyping methods we used a novel evaluation approach applied to exome sequencing data of more than 430 individuals. Several previous comparisons determined accuracy by comparing the estimated lengths of repeats to a known truth determined from either simulations or alternative assays such as PCR. Here we took only male individuals and looked at the heterozygosity of the calls only on the X chromosome. As there is only one X chromosome in males a method that reported only homozygous calls was defined to be more accurate than those that reported heterozygous calls.

### Dataset

We began with a dataset of 472 males from the Simons Simplex Collection. We had to remove 39 samples for a variety of reasons: 6 samples were not sequenced with a paired end approach, 3 samples had no coverage on the chromosome Y so were assumed to be females mislabelled as males, 28 samples produced an error in GangSTR and 2 samples could not be aligned to the reference genome by BWA due to a software error. The remaining 433 samples (Supplementary Table 1) were analysed with LobSTR (Gymrek et al., 2012), RepeatSeq (Highnam et al., 2013), HipSTR (Willems et al., 2017) and GangSTR (Mousavi et al., 2019) (see methods). In addition, variant calling was performed using the GATK best practices pipeline.

In brief, the FASTQ files were mapped using BWA-MEM (Li, 2013) and the same BAM files were used as the starting point for running each STR calling method. Each method requires a set of intervals that define repeats to be genotyped. To generate this, we used Tandem Repeats Finder (Benson, 1999) to locate tandem repeats in the hg19 reference genome and detected 224774 STR loci in the X chromosome. Because we are using exome data, we only analysed calls in the 6860 capture regions on the X chromosome. In total, we found 2322 STR loci overlapping the capture regions (Figure 1A) where almost 60% of loci consist of 6 bp repeat units (Figure 1B). In our full data set across the 433 individuals we have over a million STR loci for analysis.

**Figure 1.**
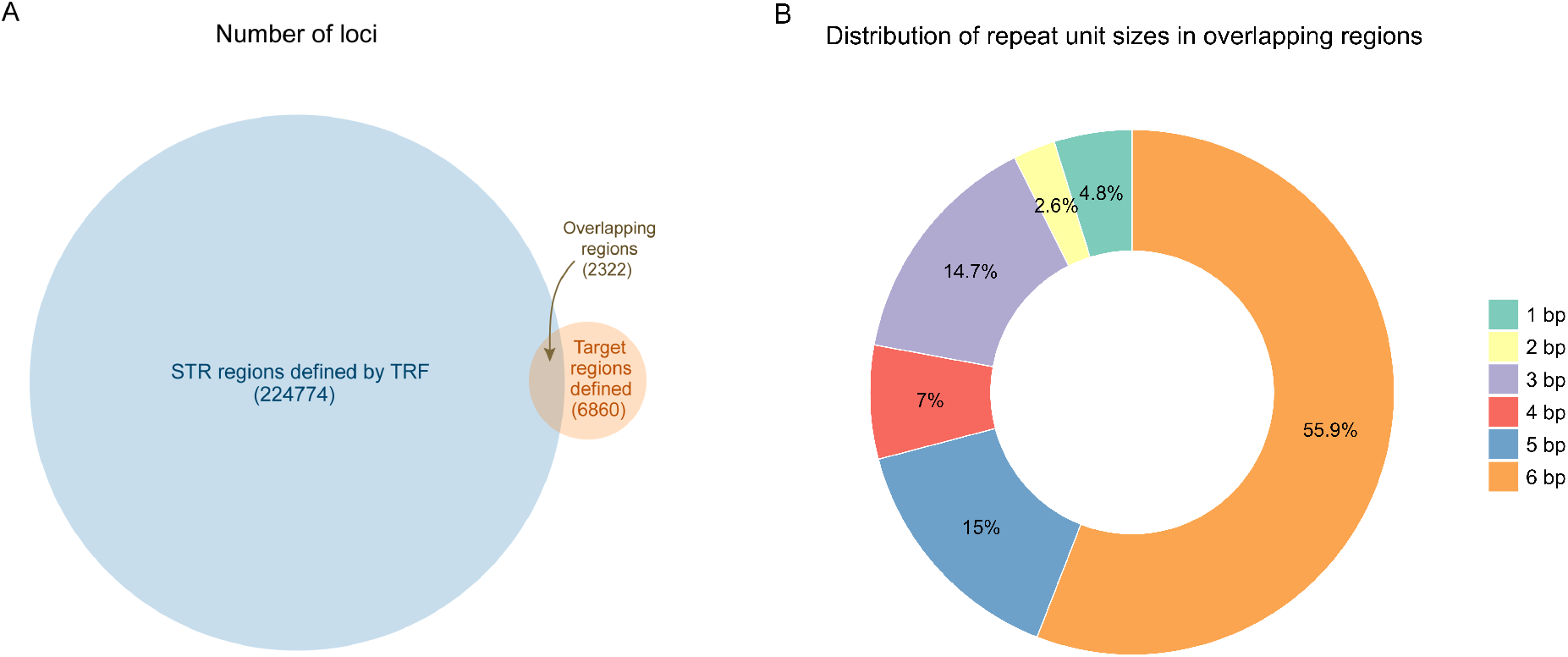
(A) Number of STR loci defined by TRF and number of regions in the capture regions of the X chromosome. Overlapping regions include all STR loci that are completely or partially overlapping the target region. Total number of STRs found in target regions is 2322. (B) Distribution of all repeat unit sizes in overlapping regions (2322).

First, we looked the ability of a method to make a call at any given locus in the capture regions. By looking at the total number for calls on the X chromosome for each method we found that LobSTR reported the highest number of loci (Figure 2A). However, the number reported for each individual was variable. Figure 2B shows the distribution of the number of reported loci per individual with the highest median number of calls by LobSTR (2015), outperforming GangSTR (1967), RepeatSeq (1847) and HipSTR (1834). GATK-HC reported a median of 11 loci per individual but rather than genotyping all loci GATK only makes a report when it deems there is a difference from the reference genome. Out of the 2322 loci we investigated there were 23 loci for which reference STR length was longer than the read length (this does not necessarily mean that the allele length is longer). Out of these 23 loci, genotypes for 6 loci were not reported by any tool in any individual. For the remaining 17 loci, all of them were reported by GangSTR in some individuals and 4 of them were genotyped by other tools as well. We also found another 13 loci were not detected by any of the tools in any of the individuals.

**Figure 2.**
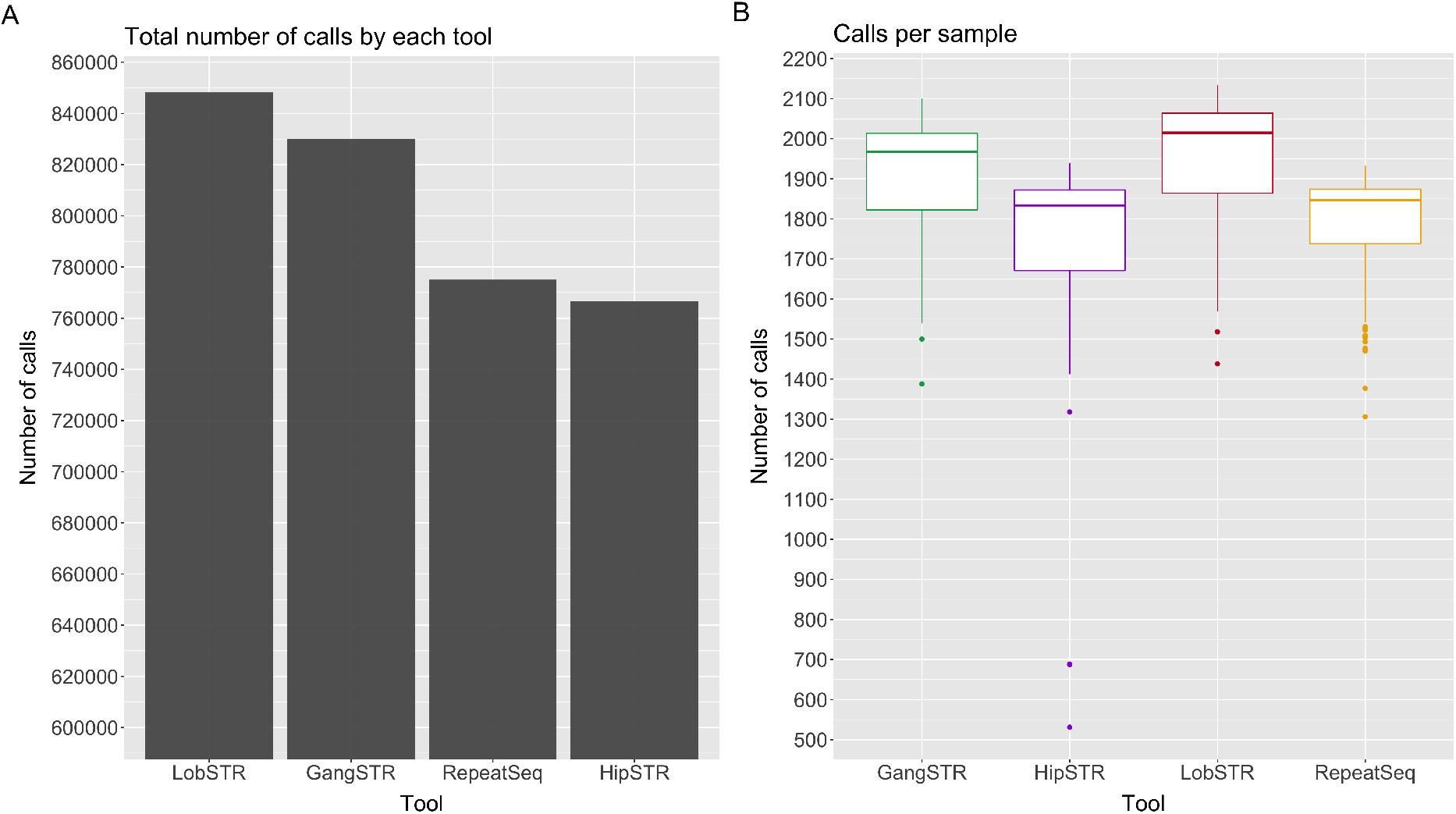
(A) Total number of calls on the X chromosome over all 433 samples made by each tool. (B) Number of calls made per sample by each tool out of a possible 2322.

GATK-HC only makes calls at positions where there is evidence that the allele is different from reference. In this dataset GATK-HC made calls in total of 346 (14.9%) different loci and 21.4% of these were heterozygous giving a minimum overall heterozygous rate of 3.1% assuming all uncalled positions are homozygous. No call either means the allele is homozygous reference or there is not enough data to make a call. This is one reason why specialised STR callers are better suited for genotyping STR loci.

To determine the genotyping accuracy of the four specialised STR callers, we first looked at the overall percentage heterozygous calls to estimate the error rate for each method. Overall, RepeatSeq had the lowest median error rate with 8675 (1.09%) of its calls being heterozygous. Next was HipSTR with 19459 (2.23%), LobSTR 27410 (2.96%) and GangSTR 33204 (3.29%). Again, error rates were variable across individual samples ranging between 0% and 47.3%. RepeatSeq, HipSTR and LobSTR are generally consistent with 3 samples outliers with respect to error rate while GangSTR has higher variability in error rates across samples (Supplementary Figure 2). Interestingly, for these three samples the heterozygous percentage increased for LobSTR after the strict filtering. All tools except RepeatSeq recommend filtering the outputs based on quality metrics of the calls (see Methods for details on filtering parameters that were used). Once the recommended filters are applied, we found that the performance of GangSTR improved by 2.76% to 0.53%, HipSTR by 2.14% to 0.09% and LobSTR by 1.32% to 1.64% (Figure 3). However, the median number of calls per sample dropped for LobSTR by 201, HipSTR by 1462 loci and GangSTR by 1512.

**Figure 3.**
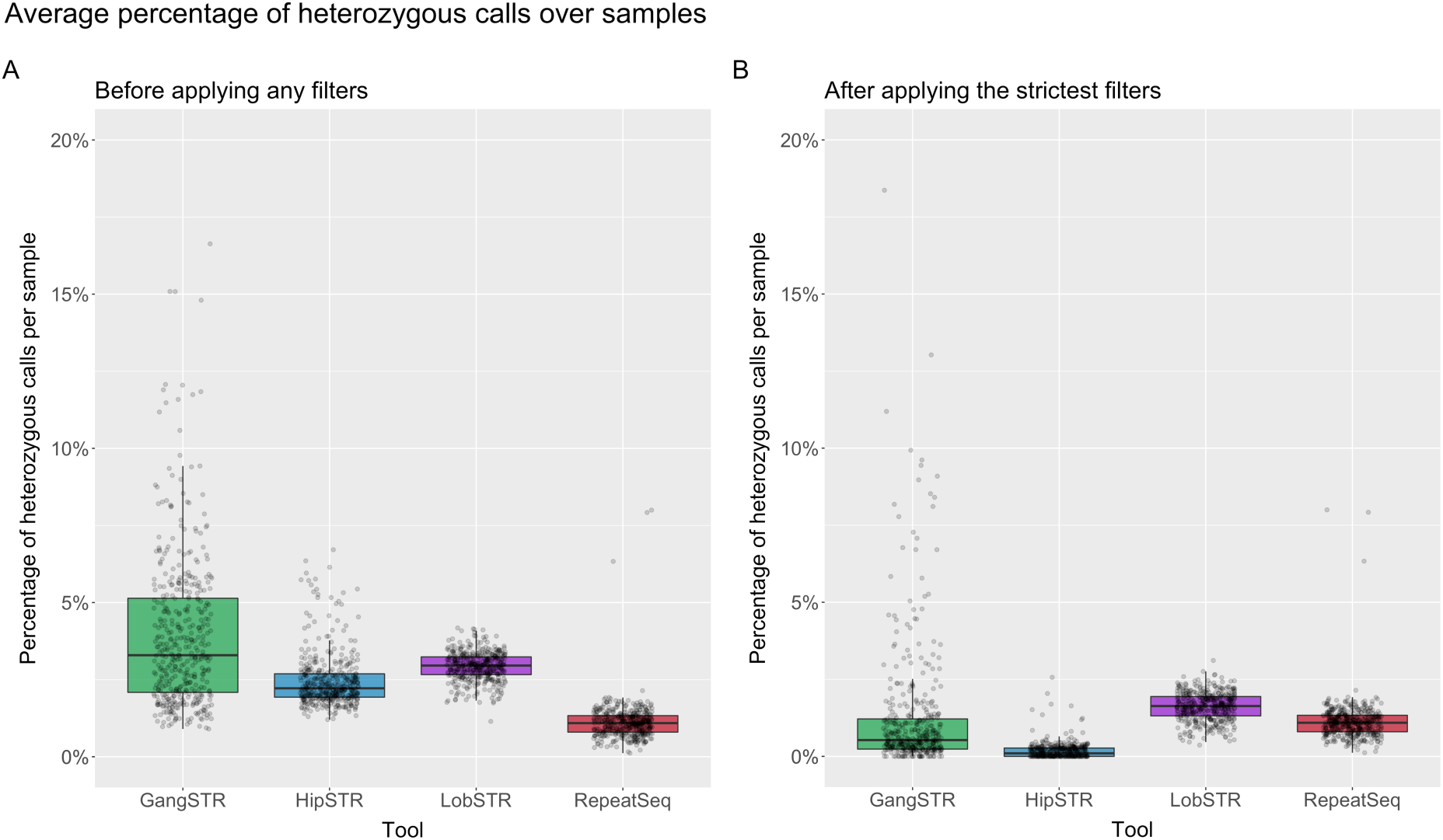
(A) Percentage of heterozygous calls over all samples (each dot is a sample) - no filters applied. (B) Percentage of heterozygous calls over all samples after applied the strictest recommended filters for the tools. Since no filters were recommended for RepeatSeq then it has the same values on both plots.

The recommended filters for each tool were different (for instance, minimum coverage of 100 for HipSTR and 50 for GangSTR) and therefore we next decided to analyse the effect of these filtering parameters separately.

### Effects of repeat unit and coverage on accuracy

We investigated the number of heterozygous calls as a function of the repeat unit length ranging from 1 to 6 bp. We found that all tools exhibit high error rates for 1 bp repeats, which is not surprising as it is difficult to genotype homopolymers due to higher rates of polymerase slippage events. More surprisingly, 2 bp repeats were poorly genotyped by HipSTR, LobSTR and GangSTR and the best results were obtained with RepeatSeq. All other repeat unit lengths produced much more accurate genotypes.

To investigate the effect of coverage and quality scores on results we applied call-level filters on our data according to developers’ recommendations to get two different datasets: low filtering where we included all suggested filters except coverage and quality scores and strict filters where we also included filters for coverage and quality scores. Then, we looked at the effects of coverage and plotted the error rate as well as percentage of remaining number of calls as a function of the minimum number of reads supporting the call. We expected the error rate to drop as coverage increased and this was the case for 3, 4, 5 and 6 bp repeat units. However, mono- and dinucleotide repeats did not follow a consistent pattern and the pattern was different between tools. While dinucleotide repeats showed a trend towards lower error rate with increasing coverage for RepeatSeq and LobSTR the trend increased for GangSTR (Figure 4).

**Figure 4.**
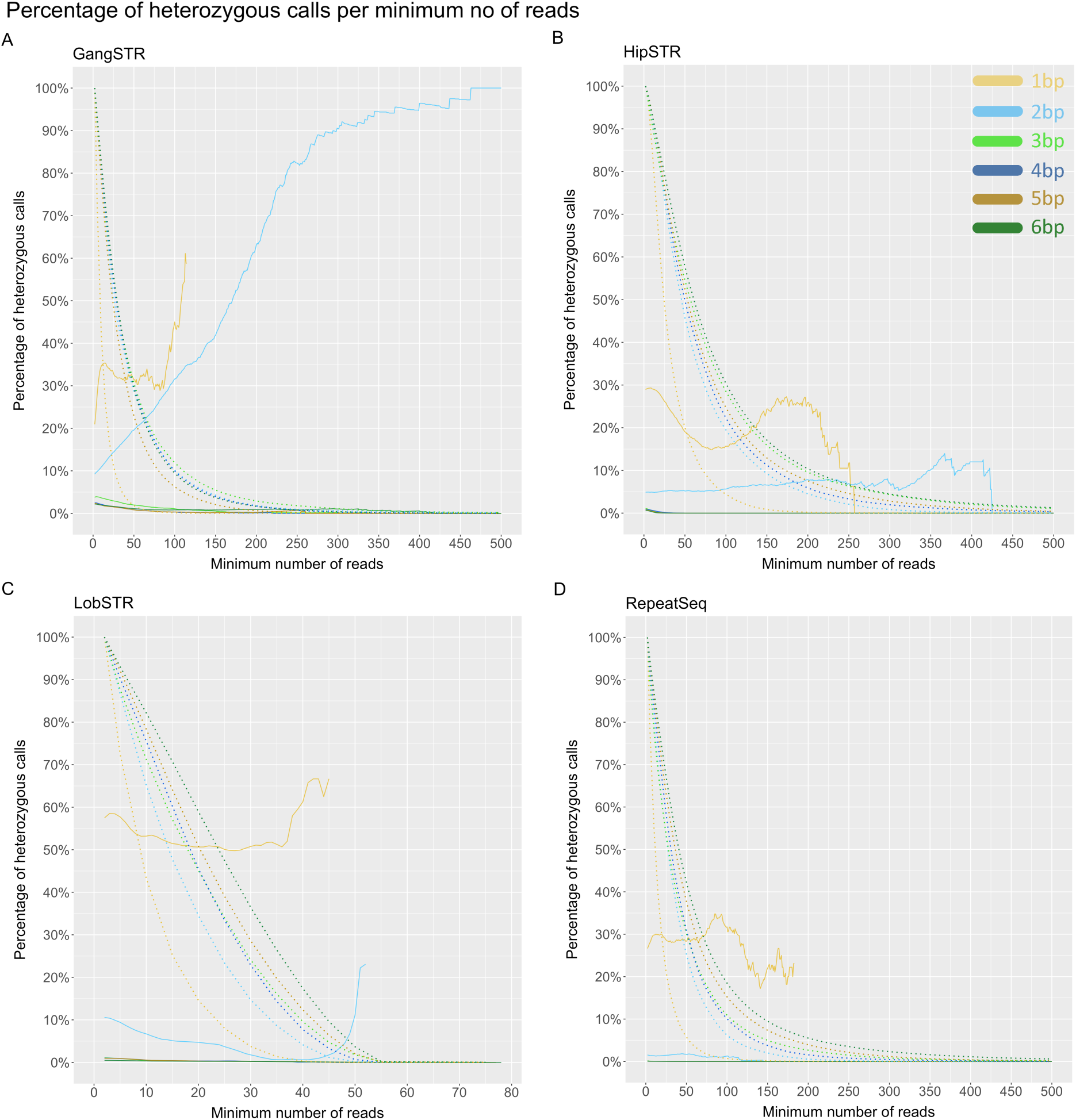
Percentage of heterozygous calls as a function of minimum number of reads. (A) GangSTR, (B) HipSTR, (C) LobSTR, (D) RepeatSeq. Solid line shows the percentage of heterozygous calls as a function of minimum number of reads. Dotted line represents the percentage of remaining calls as a function of minimum number of reads. Y-axis limited to 500 reads. Heterozygous calls are represented in percentages, but the total number of calls is different for each tool, where 100% is 796775 calls for GangSTR, 757432 calls for HipSTR, 848252 calls for LobSTR and 775030 calls for RepeatSeq.

Two call-level filters are recommended for GangSTR: level 1 that requires filtering out calls that have less than 20 reads and level 2 filters that requires at least 50 reads to support a call. Filtering out all calls with coverage of less than 20 reads brings the heterozygous error rate for the 3-6 bp repeats to between 1.6 and 3.1% at this minimum coverage. Filtering out calls with less than 50 reads improves the error rate even further to 0.83-1.97%. However, by filtering out calls with coverage less than 20 we lose on average 39.4% of 2-6 bp repeats data with a median of 1120 loci reported per sample. By filtering out calls with coverage less than 50 we lose on average 71.7% of 2-6 bp repeats data and have a median of 464 loci reported. Interestingly we see an increase of error rate as the function of coverage genotyping 2 bp STRs (Figure 4A). Unlike the other tools where heterozygous percentage decreases when coverage increases and remains relatively steady, GangSTR is not so consistent and fluctuates around 1%.

HipSTR excludes calls that have coverage less than 100 by default unless specified otherwise. This coverage gives high accuracy but also filters out 75.9% of data. We investigated reducing the minimum coverage and found HipSTR has excellent accuracy even with minimum coverage of 25: 0.02-0.04% heterozygous rate for 3, 5 and 6 bp and 0.16% for 4 bp repeats (Figure 4B). In addition, a minimum coverage of 50 improves results even more and the maximum error rate for these repeats is 0.01% at this minimum coverage (median of 899 calls per sample). HipSTR also struggles to genotype 2 bp repeats, having a heterozygous error rate around 5.2% for coverage of both 25 and 50. Therefore, by decreasing the filtering parameters for coverage to only exclude the calls less than 25x coverage we retain on average 76.6% of data for 2-6 bp repeat units with a moderate call rate (median of 1369 calls per sample).

LobSTR recommends to filter out calls with coverage less than 5 which gives us heterozygous percentage for 3-6 bp repeat units at this minimum coverage 0.44-0.90% per repeat unit length. This filters out on average 10.0% of calls for 2-6 bp repeats, giving a median of 1840 reported alleles per sample. The accuracy for these repeats increases as a function of coverage and considering the amount of calls which are filtered out, a minimum coverage of 5-10 might be the best compromise. By filtering out calls less than coverage of 10 the heterozygous error rate at this minimum coverage is 0.55% 4 bp and 5 bp repeats, 0.48% for 3 bp repeats and 0.36% for 6 bp repeats, while still retaining 74.5% of calls on average for 2-6 bp repeats (median of 1555 calls per sample). Increasing the minimum coverage to 20 reads would improve our results by a further 0.2% but also filter out significantly more calls. For dinucleotide repeats accuracy seems to get better with higher coverage and reaches 1% error rate when the minimum coverage is 33, however, the error rate increases again when coverage increases over 42 (Figure 4C).

Authors of RepeatSeq do not recommend any additional filtering and we found high accuracy even at low coverage. Filtering out all calls less than coverage of 5 will result in error rate for 3-6 bp repeat units 0.04-0.08% and only an average 7.1% of 2-6 bp repeat calls filtered out leaving an average of 1728 reported calls per sample. Among all tools, RepeatSeq shows the best accuracy for dinucleotides, having an error rate no more than 1.81% (Figure 4D).

As mentioned, we saw unusually high error rate for dinucleotide repeats in nearly all tools so we examined these repeats in more detail. Curiously, we found that in GangSTR and LobSTR the unusually high error rate of 2 bp repeats due to AC/TG repeats while other repeat units do not exhibit these characteristics (Figure 5). The same pattern was observed for RepeatSeq but at lower error rates. This information is not easily available for HipSTR as it does not report the repeat unit.

**Figure 5.**
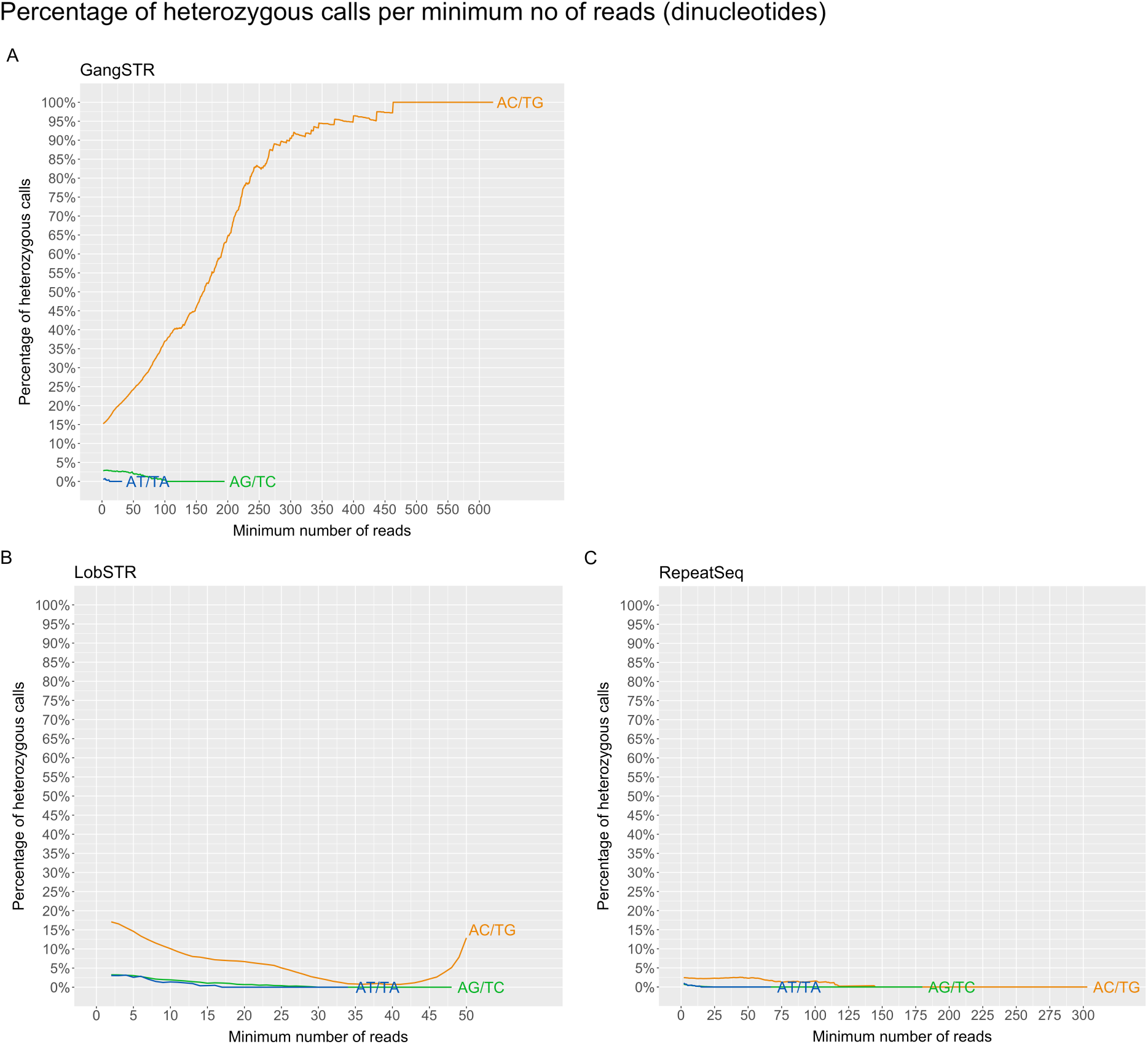
Percentage of heterozygous calls as a function of minimum number of reads for dinucleotide repeats, divided by the three possible repeat unit sequences. (A) GangSTR, (B) LobSTR, (C) RepeatSeq.

### Effects of repeat unit and quality scores on accuracy

Besides coverage, the second parameter which is commonly used for filtering in all tools is the call quality or quality score produced by each algorithm. We next investigated the effects of quality scores on the accuracy by relaxing the coverage filtering and looking at the quality scores in different bins across the score range (Figure 6). GangSTR’s level 2 filters recommends to filter out calls which have quality less than 0.9. We see on Figure 6A that the heterozygous error rate fluctuates at lower scores and starts to decrease from quality scores above 0.6 for all repeats except for homopolymers. We reach a 1% heterozygous error rate when excluding calls that have quality scores below 0.81 for 3 bp repeat units, 0.66 for 4 bp units, 0.77 for 5 bp units and 0.93 for 6 bp repeat units. We see the lowest heterozygous error rate at the quality score of 1.0 but we also determined a sharp drop in the number of reported genotypes after excluding calls with quality scores below 1.0 and we also lose 61.7% of 3-6 bp repeats data on average. The recommended 0.9 seems a reasonable suggestion for balancing the accuracy with how much data we will have left after the filtering. We can also see that accuracy of 1 bp repeat calls are improving with the highest quality score (1.0) and a stronger filter for these repeat unit may be appropriate.

**Figure 6.**
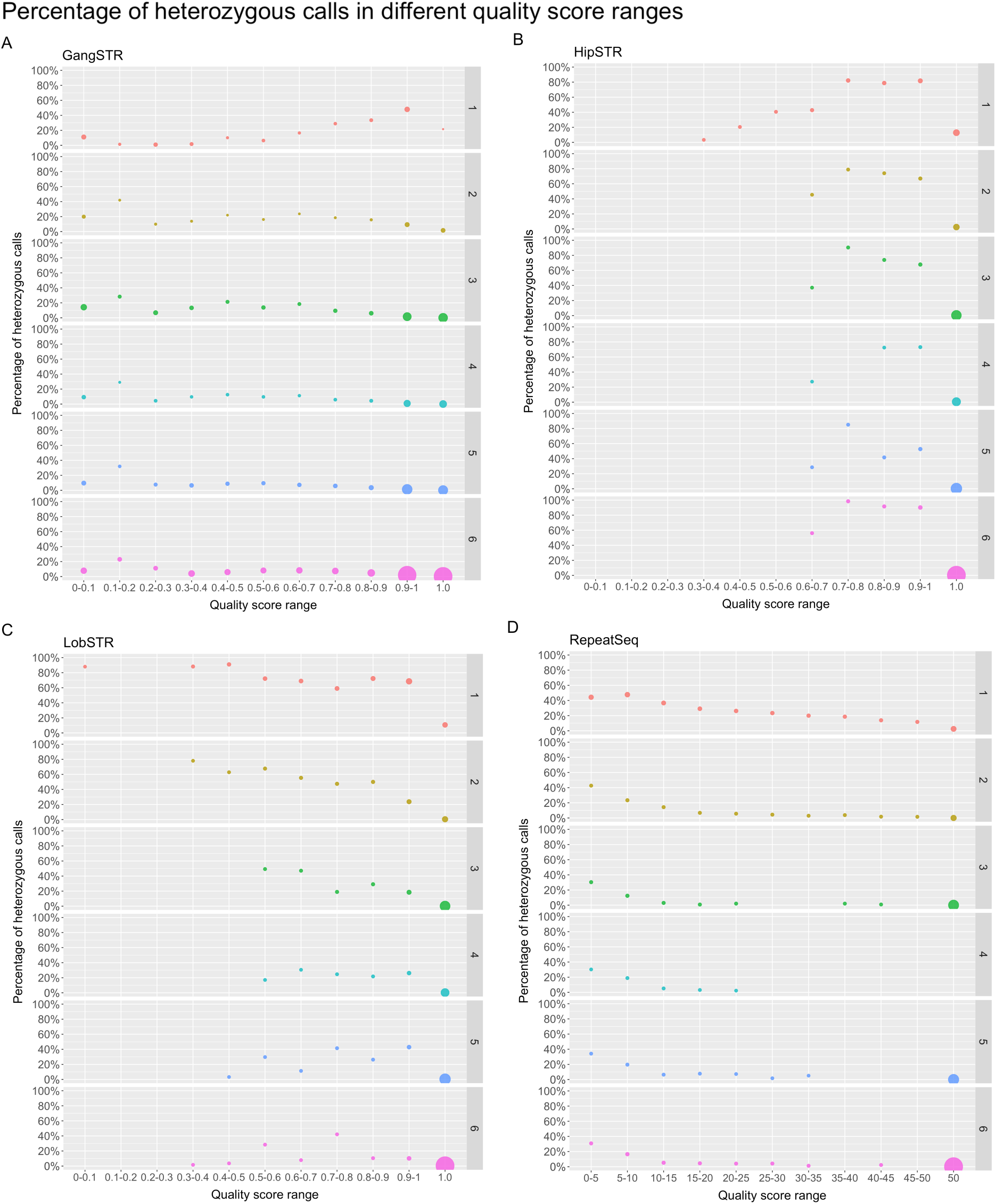
Percentage of heterozygous calls in different bins of quality score separated by each repeat unit length. (A) GangSTR, (B) HipSTR, (C) LobSTR, (D) RepeatSeq. Each bin is from marked range (inclusive) to the end value (exclusive), except the last one. Dot size represents the number of calls in that range of the repeat unit length (rows).

HipSTR, similarly to GangSTR, recommends to filter out calls with quality score less than 0.9. On Figure 6B we can see that on average the calls below the 1.0 have high heterozygous error rate and keeping only the ones with the highest quality score will improve overall accuracy. Indeed, 98.5% of calls have quality score of 1.0 and therefore we only lose a fraction of data while filtering out calls less than quality score of 1.0, which may be a good trade-off.

LobSTR recommends to filter out calls with call quality score less than 0.8. As seen on Figure 6C heterozygous error rate fluctuates but generally shows more accuracy with higher quality scores for 1-3 bp repeats, while for 4-6 bp there is no clear trend below quality score of 0.9. There is no significant improvement in overall accuracy when we remove calls with quality scores less than 0.8, or even 0.9, which might be due to the fact that, similar to HipSTR, the majority of calls (94.3%) have quality score of 1.0. We do see improvement while only leaving calls with quality of 1.0 particularly in 1-2 bp repeat units which shows utility in filtering out the least accurate calls and since the majority of data has quality score of 1.0 this filtering could be a good choice, as also suggested for HipSTR (see Supplementary Figure 3).

RepeatSeq does not recommend any filtering and reports quality scores on a Phred scale (Figure 6D). We determined that filtering out calls with Phred quality score of 10 or less improves the accuracy of all repeat units. Accuracy of genotyping mono- and dinucleotide repeats continues to improve as a function of Phred scores while the best accuracy is observed at the highest quality score. On the flip side, the number of calls also decreases, and at the highest Phred scores, we are left with 31.6% of homopolymers and 87.8% of dinucleotide repeats. However, it only filters out 1.2% of 3-6 bp repeats data. Overall, filtering data based on the quality scores may be reasonable if looking at mono- or dinucleotide repeats and accuracy is an important factor.

### Accuracy of GangSTR by looking at only the enclosing class of reads

LobSTR, HipSTR and RepeatSeq are using types of reads where the STR region has to be completely in the read. However, GangSTR uses more classes of reads that may give rise false positives. Therefore to make a more direct comparison we decided to look at GangSTR results where we filtered out all other classes of reads besides the enclosing ones, marked here as GangSTR (enc.). Compared to the previous GangSTR results we now see a lower error rates for all repeat units as well as no substantial fluctuation on higher coverage that was apparent previously on Figure 4A. At the coverage of 20, GangSTR (enc.) has heterozygous error rate 0.53-0.99% for 3-6 bp repeats while 58.6% of 2-6 bp repeats data is filtered out, and with a median of 699 calls per sample (Figure 7A). When we increase the minimum number of reads to 50 we can see even further improvement, 0.01-0.27% error rate for 3-6 bp repeats, however, also filtering out 86.3% of 2-6 bp repeats data (median of 152 calls per sample).

**Figure 7.**
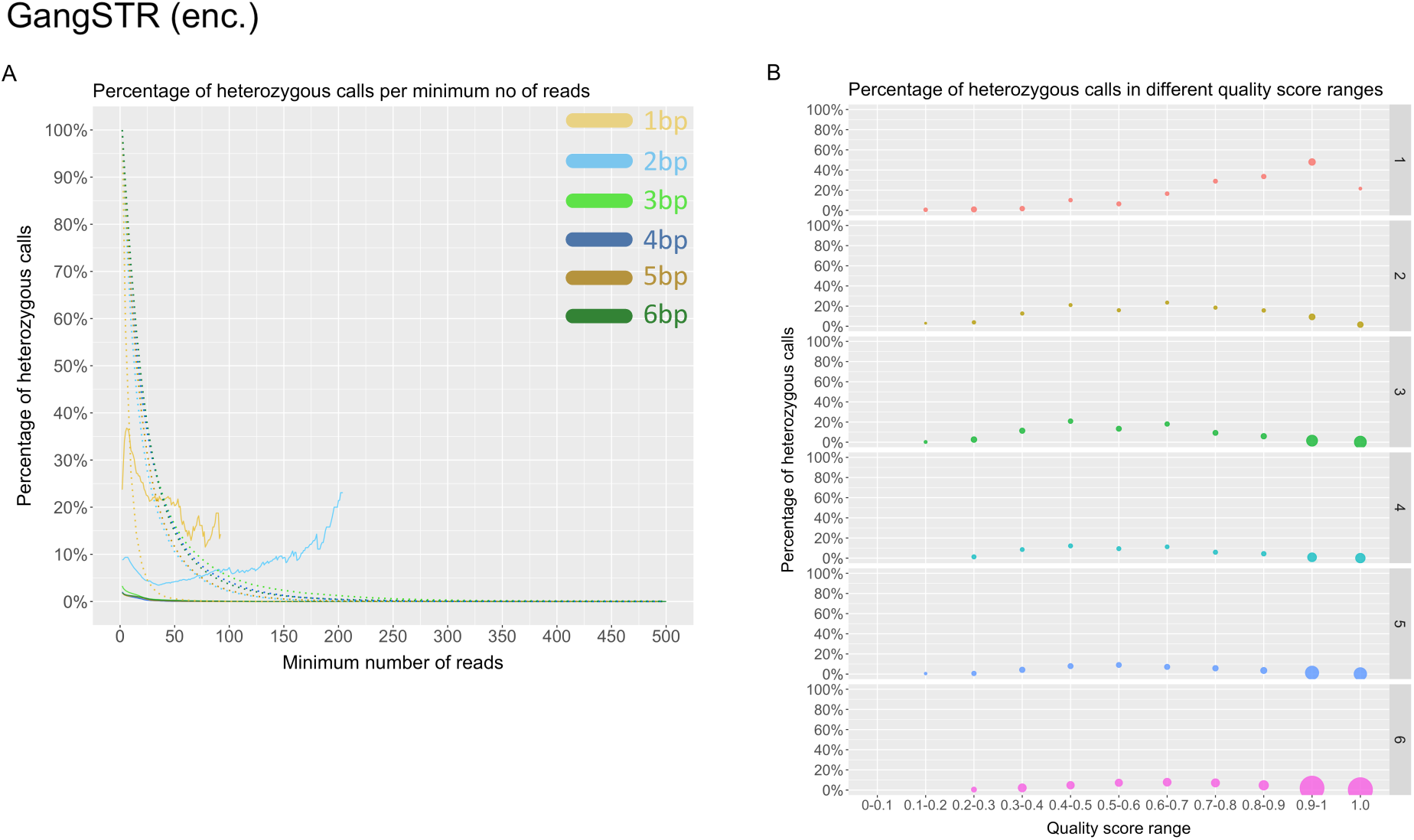
(A) Percentage of heterozygous calls as a function of minimum number of reads. Solid line represents the percentage of heterozygous calls as a function of minimum number of reads or quality score and dotted line the percentage of remaining calls as a function of minimum number of reads or quality score. (B) Percentage of heterozygous calls in different ranges of quality scores. Each bin is from marked range (inclusive) to the end value (exclusive), except the last one. Dot size represents the number of calls in that range of the repeat unit length (rows).

We also looked at the dinucleotides separately (see Supplementary Figure 4) and found that it is considerably lower compared to results of GangSTR. However, we determined that AC/TG repeats still have a higher error rate compared to other repeat units (Figure 4A and Figure 7A). Results of quality scores are quite similar to the GangSTR’s where we see improvements at high quality scores (Figure 7B).

### Run time of tools

Finally, we benchmarked all tools to determine their average running time. We ran each tool genome wide by using 1, 2, 4 and 8 cores and determined that only RepeatSeq is supporting multithreading, which allows the tool to run faster when utilising more processor cores. In particular, we saw that for samples that have coverage on target regions around 90x, the average running time on 1 core for LobSTR was 9 min 31 sec giving it a clear advantage compared with, HipSTR at 1 h 22 min, GangSTR at 3 h 4 min and RepeatSeq 2 h 58 min. By utilizing more cores, LobSTR’s and GangSTR’s and HipSTR’s running time remained the same, while RepeatSeq’s time improved.

## Discussion

We used a novel way of evaluating the error in short tandem repeats genotyping methods where we analysed STR calls on the X chromosome of male samples. Because this is a hemizygous chromosome we determined a relative error rate as the rate of heterozygous genotypes. We performed this evaluation on a human exome dataset of 433 samples resulting in the evaluation of more than a million STR loci. Exome sequencing is widely used, but the PCR step in library preparation causes a challenge to STR genotyping tools due to the interference of stutter noise. This is the first independent evaluation of these STR genotyping tools that we are aware of.

Many of the tools do not claim to be able to accurately genotype homopolymers and we found that indeed all tools had difficulty with these repeats resulting in a high error rate. There was also no clear correlation between minimum coverage and accuracy of genotyping homopolymers, but using the highest quality scores did improve the accuracy. Interestingly, most tools produced high error rates for genotyping dinucleotides as well, which we later found to be mainly caused by AC/TG repeat units. One who analyses dinucleotide repeats with these tools should be aware of the differences in accuracy of genotyping different repeat units and carefully interpret the results of AC/TG repeat units. Repeat units with length of 3-6 bp were all relatively accurate and similar across tools with only minor differences. However, genotyping was slightly less accurate for 3 bp length repeats in low coverage and low quality scores, but differences were reduced with proper filtering. We found that LobSTR was able to report the highest number of genotypes at a heterozygous error rate of less than 1%.

There are certain filtering parameters suggested for each tool and we examined the effects of coverage and quality scores across all tools. However, some tools have further parameters that could be explored that were not part of our investigation. In general, we found that higher quality scores increased the accuracy of results at the cost of losing some potentially accurate calls. The relationship with coverage was more complex but some coverage filtering improved results for all tools. Which parameters to use depends on the aim of the analysis. For example, more calls may be desirable to begin a screen, or more accuracy may be desirable if selecting potential disease associated loci. When one does an exploratory analysis to find potential loci of interest that can be followed up with alternative methods then lowering filtering parameters for coverage and quality scores for certain tools could be a good approach as it leaves us with larger portion of data. We found that even in exome data we can use these tools to genotype tens of thousands of loci.

Unlike the other tools we used in this analysis, GangSTR is utilising four different types of reads which can help to pick up the locus other tools cannot (such as those longer than the read length). However, these can also produce genotyping errors. In our analysis, we first looked at GangSTR results that included all four classes of reads and then we excluded all calls where only spanning or bounding class of reads were present, as suggested by the tool authors. This filtering increased the genotyping accuracy of the tool (we also looked at the results where we skipped this filtering parameter but this did not improve results). Still, compared to other tools, GangSTR showed higher error rate. Finally, we decided to look at only enclosing class of reads such as the other tools are doing and determined around three times lower error rate at 20x and bigger gains on higher coverage. On the other hand, that change will result in losing the ability to genotype alleles longer than the read length that is GangSTR’s important addition. We also found that HipSTR has a very high accuracy for 3-6 bp repeats when coverage is at least 50x. Excellent accuracy was also found in RepeatSeq in very low coverage and genotyping dinucleotides were the most accurate among the tools. In addition, RepeatSeq is the only tool that supports multithreading and therefore can run faster by allocating more cores.

Here we have presented one way of performing an evaluation and this approach does not look at accuracy of the estimated allele length which is a limitation of the study. In addition, it is difficult to rule out a bias towards tools that default to genotyping an allele as a homozygous reference by the software. Our comparisons were specifically analysing the exome dataset that was PCR amplified where a tools’ noise model may play an important role. Therefore, tools may perform differently when we will analyse PCR−free WGS dataset.

In conclusion, all these tools are built to genotype STRs but have different strengths and weaknesses. Based on our analysis there is no clear overall winner. RepeatSeq and HipSTR are the best when considering genotyping error rate even with low coverage. On the other hand, GangSTR has an advantage because it is the only tool among them that can call alleles longer than the read length but shows higher error rate unless looking at only the enclosed class of reads, which in turn would lose the GangSTR’s advantage of picking up long genotypes. In addition, GangSTR is the newest tool and so comes with reference files for different reference builds that are periodically updated according to the tool’s webpage. The correct choice of a tool and the subsequent filtering depends on the aim of the analysis, and might be influenced by available hardware resources and time limit for running tools.

## Methods

In order to compare all STR tools, we ran each one of them on the same dataset. We used the data from the publicly available Simons Simplex Collection (SSC) for our analysis.

Samples’ genomic DNA was extracted from whole blood. Exomes were captured with NimbleGen EZ Exome v2.0 (Roche Nimblegen, Inc., Madison, WI) reagents and sequenced using Illumina (San Diego, CA) GAIIx (N = 271) or HiSeq 2000 (N = 162) at the Yale University School of Medicine.

All SRA files were downloaded from NCBI and converted to FASTQ files. In total there were 238 families where only males were selected for our analysis to avoid heterozygous sites in the X chromosome, assuming that any multiallelic STR calls should be a result of PCR and/or sequencing error. Male samples were determined by using metadata of samples (472 samples) and quality controlled by looking at the coverage on X and Y chromosome. Results of the analysis lead to exclusion of 3 samples as they had no coverage on Y chromosome. Out of the remaining 469 samples, we excluded 6 single-end read sequenced files as well as 28 paired-end read sequenced samples that did not work on GangSTR, and 2 additional samples that had issues with mapping, which left us with 433 samples in total (Supplementary Table 1).

All computational steps (tools and parameters used) are described in this section.

### SRA to FASTQ conversion

All whole-exome sequencing SRA files in Simons Simplex families (study’s identifier: SRP010920) were downloaded from NCBI and converted to FASTQ files by using *fastq-dump* (https://ncbi.github.io/sra-tools/fastq-dump.html):

~~~
fastq-dump \
       --gzip \
       --skip-technical \
       --readids \
       --read-filter pass \
       --split-3 \
       --dumpbase \
       --clip \
       FILE.SRA
~~~

### Defining STR regions

The human reference sequence hg19 (February 2009 assembly) was downloaded from UCSC (http://hgdownload.soe.ucsc.edu/goldenPath/hg19/bigZips/hg19.fa.gz) and “hg19.fa” file was created by and indexed with Samtools v1.10 (Li et al., 2009) which was downloaded from http://www.htslib.org website:

~~~
cat *.fa > hg19.fa
samtools faidx hg19.fa
~~~

Creating FASTA sequence dictionary file for GATK analysis:

~~~
gatk CreateSequenceDictionary
       -R hg19.fa
       -O hg19.dict
~~~

Tandem Repeats Finder v4.09 (Benson, 1999) was downloaded from https://tandem.bu.edu/trf/trf.html and used to find STRs (1-6 bp repeat unit length) in hg19 reference genome by using the following command and parameters:

~~~
./trf409.linux64 hg19.fa 2 7 7 80 10 24 6 -h
~~~

A custom-made Python script named trf2bed (https://gitlab.com/andreassh/trf2bed) was used to extract data from TRF output file to generate BED regions file for LobSTR, GangSTR, HipSTR, RepeatSeq and GATK.

~~~
python3 trf2bed.py \
        --dat hg19.fa.2.7.7.80.10.24.6.dat \
        --bed hg19.fa.2.7.7.80.10.24.6_$TOOL.bed \
        --tool $TOOL
~~~

### FASTQ alignment and calculating BAM coverage

Reads from FASTQ files were aligned to hg19 reference genome using BWA-MEM v0.7.17 that was downloaded from https://sourceforge.net/projects/bio-bwa/files/ and aligned BAM files were merged and indexed by using Samtools v1.10 (Li et al., 2009)

~~~
bwa mem -M -t 8 -R “@RG\\tID:$id\\tPL:$PLATFORM\\tPU:$BARCODE\\tSM:$SAMPLE\\tLB:$LIBRARY”
hg19.fa $INPUT_FILE1.fastq $INPUT_FILE2.fastq | samtools sort -@hg19.fa -o $OUTPUT_FILE.bam -

samtools merge $OUTPUT_FILE.merge.bam $INPUT_FILE1.bam $INPUT_FILE2.bam
samtools index $INPUT_FILE.merge.bam
~~~

To follow the best practices of GATK duplicate reads were removed:

~~~
gatk MarkDuplicatesSpark \
        INPUT=$INPUT_FILE.merge.bam \
        OUTPUT=$OUTPUT_FILE.bam \
        --remove-sequencing-duplicates \
        --create-output-bam-index
~~~

Coverage of BAM files on target regions was found with MosDepth v0.2.4 tool that was downloaded from https://github.com/brentp/mosdepth/ followed by calculating the median and average coverage:

~~~
mosdepth \
        -n \
        --fast-mode \
        --by $TARGET_REGIONS.bed \
        $OUPUT_FILE.coverage \
        $INPUT_FILE.merge.bam
gunzip -c $INPUT_FILE.regions.bed.gz | sort -n -k 5 | awk
‘{a[NR]=\$5}END{print(NR%2==1)?a[int(NR/2)+1]: (a[NR/2]+a[NR/2+1])/2}’ > $OUTPUT_FILE.avgcov

gunzip -c $INPUT_FILE.regions.bed.gz | sort -n -k 5 | awk ‘{ sum += \$5; n++ } END { if (n > 0) print sum / n; }’ > $OUTPUT_FILE.medcov
~~~

### Calling STRs and genotyping

GangSTR v2.4 (Mousavi et al., 2019) was downloaded from https://github.com/gymreklab/GangSTR and executed with the following parameters:

~~~
GangSTR \
       --bam $INPUT_FILE.merge.bam \
       --ref hg19.fa \
       --regions hg19.fa.2.7.7.80.10.24.6_gangstr.bed \
       --out $OUTPUT.vcf \
       --nonuniform \
       --coverage X*

*\* where X = mean coverage for the particular sample that was calculated by MosDepth tool as described previously*.
~~~

Strict filtering was done as recommended by the developer by using dumpSTR that is part of STRTools package (https://github.com/gymreklab/STRTools):

~~~
dumpSTR \
       --vcf $INPUT_FILE.vcf \
       --out $OUTPUT_FILE \
       --filter-spanbound-only \
       --filter-badCI \
       --max-call-DP 1000 \
       --min-call-DP 50 \
       --min-call-Q 0.9
~~~

Since we were looking the relationship between coverage and quality scores, and genotyping accuracy separately, we did additional filtering (partial filtering) where we discarded the filtering on calls with low coverage or low-quality scores:

~~~
dumpSTR \
       --vcf $INPUT_FILE.vcf \
       --out $OUTPUT_FILE \
       --filter-spanbound-only \
       --filter-badCI \
       --max-call-DP 1000
~~~

RepeatSeq v0.8.2 (Highnam et al., 2013) was downloaded from https://github.com/adaptivegenome/repeatseq and executed with following parameters:

~~~
repeatseq \
       $INPUT_FILE.bam \
       hg19.fa \
       hg19.fa.2.7.7.80.10.24.6_repeatseq.bed
~~~

LobSTR v4.0.6 (Gymrek et al., 2012) was downloaded from http://lobstr.teamerlich.org and custom lobSTR reference was made using lobstr_index.py and GetSTRInfo.py scripts as follows:

~~~
python ./lobstr/scripts/lobstr_index.py
       --str hg19.fa.2.7.7.80.10.24.6_lobstr.bed \
       --ref hg19.fa \
       --out ./lobstr/hg19_custom/
python ./lobstr/scripts/GetSTRInfo.py \
       hg19.fa.2.7.7.80.10.24.6_lobstr.bed hg19.fa >
       ./lobstr/hg19_custom/lobstr_hg19_custom_strinfo.tab
~~~

LobSTR’s allelotype was used to call STRs and it was running with default parameters, with and without the the *--no-rmdup* flag:

~~~
./lobstr/bin/allelotype \
       --command classify \
       --bam $INPUT_FILE.merge.bam \
       --index-prefix ./lobstr/hg19_custom/lobstr_hg19_custom_ref/lobSTR_ \
       --strinfo ./lobstr/hg19_custom/lobstr_hg19_custom_strinfo.tab \
       --noise_model ./lobstr/share/lobSTR/models/illumina_v3.pcrfree \
       --out $OUTPUT_FILE.vcf \
       --no-rmdup
~~~

Willems and colleagues explored the effects of recommended allelotype options for lobSTR (*--filter-mapq0, --filter-clipped, --max-repeats-in-ends* and *--min-read-end-match*), but found the optimal settings for lobSTR does not include these parameters and best results are obtained with default ones, which was reported about RepeatSeq (Willems et al., 2017) and therefore we decided to run both tools with the default parameters.

We did the strict filtering with the lobSTR’s filtering tool, based on the author’s recommendations for a whole genome data:

~~~
python ./lobstr/share/lobSTR/scripts/lobSTR_filter_vcf.py \
       --vcf $INPUT_FILE.vcf > $OUTPUT_FILE.vcf \
       -loc-cov 5 \
       --loc-log-score 0.8 \
       --loc-call-rate 0.8 \
       --loc-max-ref-length 80 \
       --call-cov 5 \
       --call-log-score 0.8 \
       --call-dist-end 20
~~~

And the partial filtering:

~~~
python ./lobstr/share/lobSTR/scripts/lobSTR_filter_vcf.py \
       --vcf $INPUT_FILE.vcf > $OUTPUT_FILE.vcf \
       --loc-call-rate 0.8 \
       --loc-max-ref-length 80 \
       --call-dist-end 20
~~~

HipSTR v0.6.2 (Willems et al., 2017) was downloaded from https://github.com/tfwillems/HipSTR and executed with following parameters:

~~~
HipSTR \
       --min-reads 2 \
       --def-stutter-model \
       --fasta hg19.fa \
       --regions hg19_2.7.7.80.10.24.6_hipstr.bed \
       --str-vcf $OUTPUT_FILE.vcf.gz \
       --bams $INPUT_FILE.merge.bam
~~~

Strict filtering was done according to the developer’s recommendations:

~~~
python ./HipSTR/scripts/filter_vcf.py \
       --vcf $INPUT_FILE.vcf \
       --min-call-qual 0.9 \
       --max-call-flank-indel 0.15 \
       --max-call-stutter 0.15 \
       --min-call-allele-bias -2 \
       --min-call-strand-bias -2 > $OUTPUT_FILE.vcf
~~~

Since we ran HipSTR with *--min-reads 2* parameter, we additionally filtered out all calls that had less than 100 reads, as this is the default parameter that HipSTR uses.

Partial filtering was done:

~~~
python ./HipSTR/scripts/filter_vcf.py \
       --vcf $INPUT_FILE.vcf \
       --max-call-flank-indel 0.15 \
       --max-call-stutter 0.15 \
       --min-call-allele-bias -2 \
       --min-call-strand-bias -2 > $OUTPUT_FILE.vcf
~~~

GATK v4.1.2 was downloaded from https://software.broadinstitute.org/gatk/download/ and executed with following standard parameters:

~~~
gatk HaplotypeCaller \
       --reference hg19.fa \
       --intervals hg19_2.7.7.80.10.24.6_gatk.bed \
       --genotyping-mode DISCOVERY \
       --input $INPUT_FILE.merge.bam \
       --output $OUTPUT_FILE.vcf
~~~

Data were analysed using GATK best practice guidelines (DePristo et al., 2011) up to variant calling. Variant calling was performed with the HaplotypeCaller in GATK (DePristo et al., 2011).

### Data extraction from variant calling files and analysis

A custom-made Python script was created to extract data from outputted variant calling files (VCF) of all tools ran. In particularly, only calls in the X chromosome were extracted out. In case of filtering, only calls that passed the filter were extracted out. The output file contained information about all STR loci found in the VCF file, having the following fields: sample name, locus, chromosome, start and end coordinates of the STR region, motif (repeat unit), length of motif, length of the reference, length of alleles, genotype, number of total reads, number of reads supporting the call in each class and the quality score.

The data was then analysed in R. Bioconductor’s GenomicRanges package for R (Lawrence et al., 2013) was used to find and then filter out all calls that fell outside of the target regions. All STR regions that were entirely or partially inside of the target region were included in the analysis, however, all duplicate loci were removed. When calculating heterozygous percentage per minimum number of reads or quality score bins, we only included the results when there were minimum of ten results (samples) to use and calculated the percentage again for each repeat unit length after each read or after 1/10 quality score bin.

### Running time and multithreading

To see the performance of the STR specific tools we selected 5 WES samples that had a median coverage on target regions closest to 90x (between 88.6x and 91.6x) and calculated the time each tool ran on each sample individually by using either 1, 2, 4 or 8 processor cores on the same server that has Intel(R) Xeon(R) 2.60 GHz processors and maximum of 16 GB RAM. Each test was repeated for 3 times and calculated the average time. Timing was performed with the UNIX time command.

## Declarations

## Acknowledgments

The authors are thankful to Katrina Bell for their comments and suggestions for the article.

## Author Contributions

The project was conceived and the manuscript written by A.H. and A.O. Data analysis was performed by A.H. with critical input from A.O. All authors read and approved the final manuscript.

## Data Availability

The Simons Simplex Collection dataset can be downloaded from NCBI (http://www.ncbi.nlm.nih.gov/sra). List of IDs of samples used in the analysis can be found in the Supplementary Table 1.

