## Supplementary for "Accuracy of short tandem repeats genotyping tools in whole exome sequencing data"

### Supplementary information

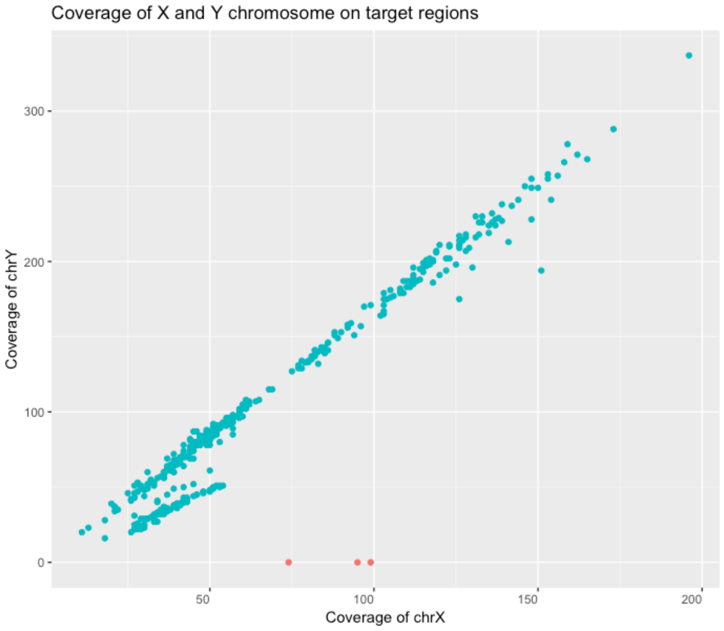


Figure 1. Coverage of X and Y chromosome of samples marked as male in the metadata.

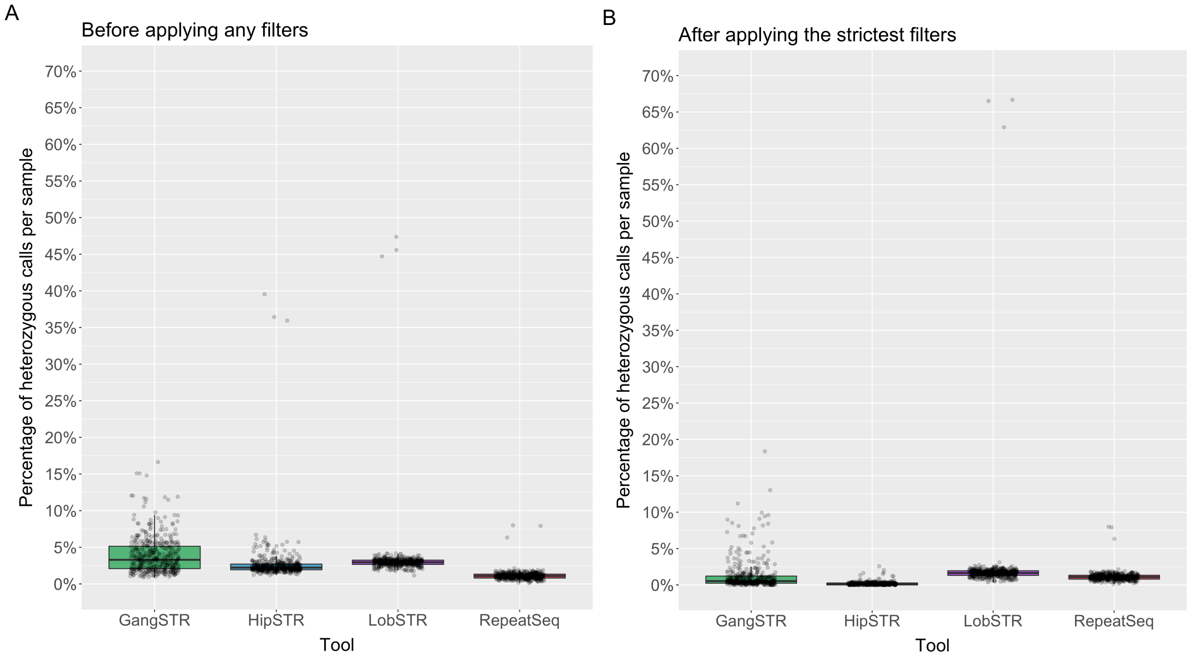


Figure 2. (A) Percentage of heterozygous calls over all samples (each dot is a sample) - no filters applied. (B) Percentage of heterozygous calls over all samples after applied the strictest recommended filters for the tools. Since no filters were recommended for RepeatSeq then it has the same values on both plots.

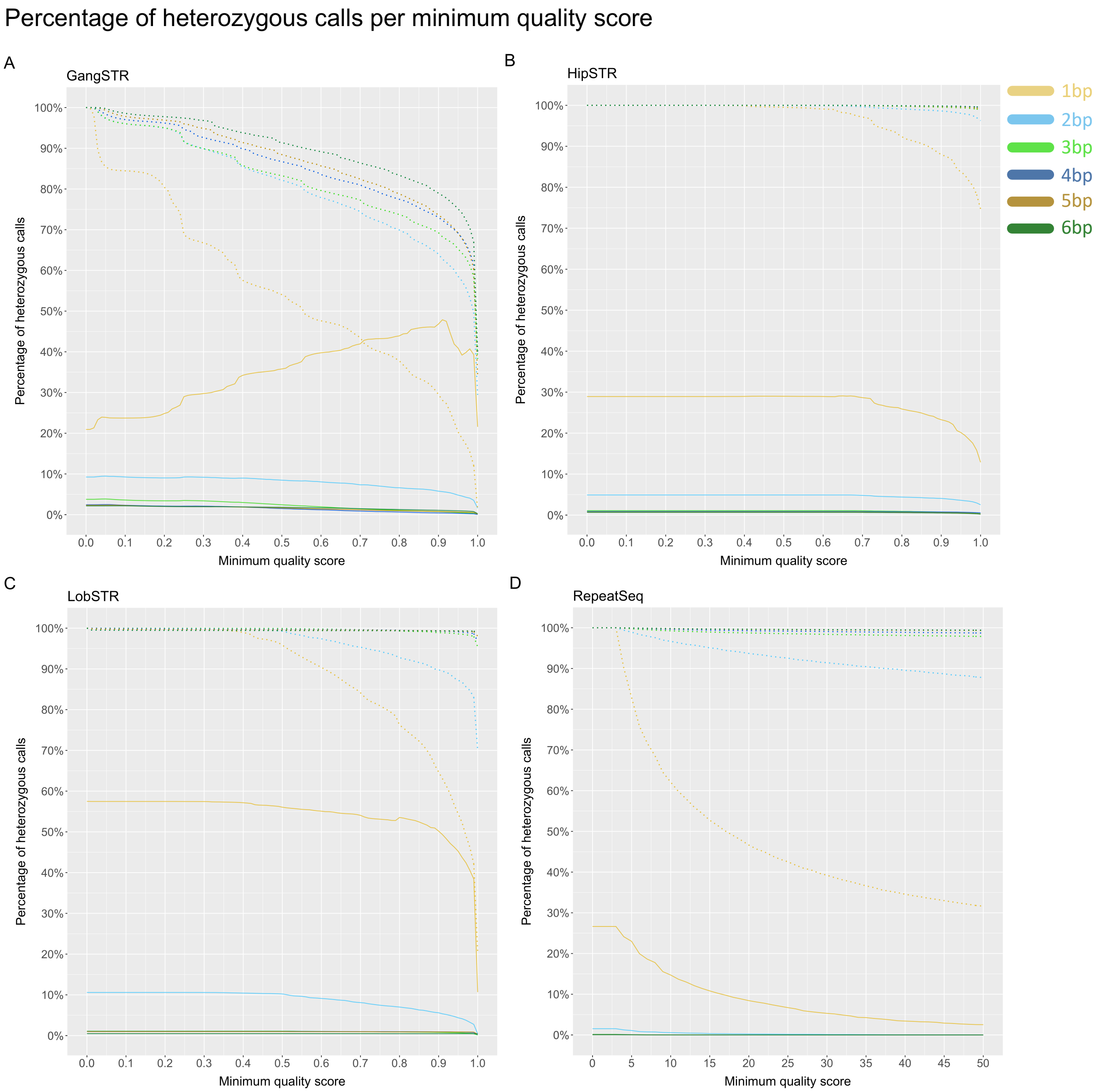


Figure 3. Percentage of heterozygous calls as a function of minimum quality score. (A) GangSTR, (B) HipSTR, (C) LobSTR, (D) RepeatSeq. Solid line represents the percentage of heterozygous calls as a function of minimum quality score and dotted line the percentage of remaining calls as a function of minimum quality score. Dotted line represents the percentage of remaining calls as a function of minimum number of reads. Heterozygous calls are represented in percentages, but the total number of calls is different for each tool, where 100% is 796775 calls for GangSTR, 757432 calls for HipSTR, 848252 calls for LobSTR and 775030 calls for RepeatSeq.

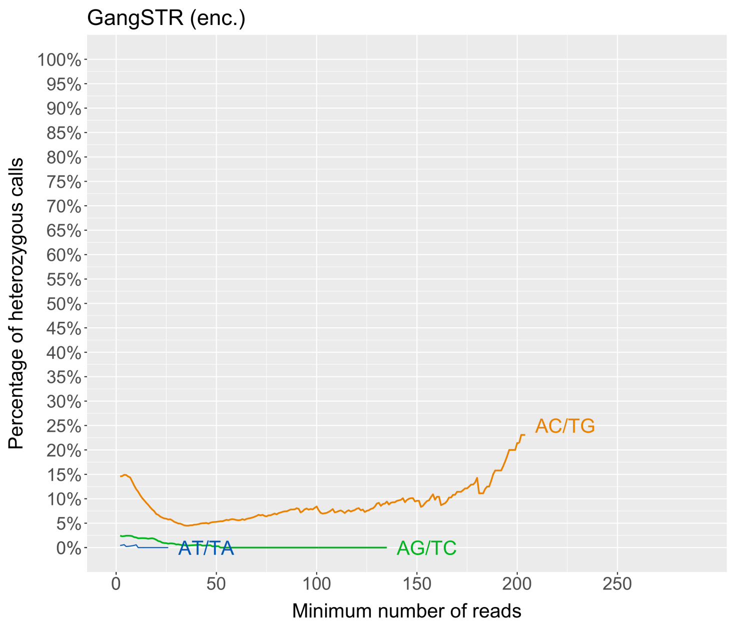

Figure 4. Percentage of heterozygous calls per minimum number of reads (dinucleotides).

Table 1. List of samples included in the analysis.

| | | | | | |
|---|---|---|---|---|---|
| SRR1272193 | SRR1272230 | SRR1272231 | SRR1272233 | SRR1272235 | SRR1272237 |
| SRR1272240 | SRR1272241 | SRR1272242 | SRR1272244 | SRR1272245 | SRR1272246 |
| SRR1272248 | SRR1272250 | SRR1272251 | SRR1272253 | SRR1272254 | SRR1272255 |
| SRR1272257 | SRR1272258 | SRR1272259 | SRR1272263 | SRR1272267 | SRR1272271 |
| SRR1272274 | SRR1272275 | SRR1272278 | SRR1272280 | SRR1272282 | SRR1272284 |
| SRR1272285 | SRR1272287 | SRR1272288 | SRR1272289 | SRR1272291 | SRR1272293 |
| SRR1272295 | SRR1272296 | SRR1301226 | SRR1301227 | SRR1301228 | SRR1301229 |
| SRR1301230 | SRR1301232 | SRR1301233 | SRR1301234 | SRR1301236 | SRR1301237 |
| SRR1301238 | SRR1301240 | SRR1301241 | SRR1301242 | SRR1301245 | SRR1301246 |
| SRR1301248 | SRR1301250 | SRR1301252 | SRR1301253 | SRR1301254 | SRR1301256 |
| SRR1301257 | SRR1301258 | SRR1301262 | SRR1301264 | SRR1301266 | SRR1301270 |
| SRR1301272 | SRR1301273 | SRR1301274 | SRR1301276 | SRR1301277 | SRR1301280 |
| SRR1301281 | SRR1301283 | SRR1301284 | SRR1301286 | SRR1301287 | SRR1301289 |
| SRR1301290 | SRR1301291 | SRR1301293 | SRR1301295 | SRR1301297 | SRR1301299 |
| SRR1301303 | SRR1301307 | SRR1301310 | SRR1301313 | SRR1301314 | SRR1301316 |
| SRR1301318 | SRR1301320 | SRR1301322 | SRR1301324 | SRR1301326 | SRR1301328 |
| SRR1301329 | SRR1301330 | SRR1301332 | SRR1301334 | SRR1301336 | SRR1301337 |
| SRR1301338 | SRR1301340 | SRR1301341 | SRR1301342 | SRR1301344 | SRR1301345 |
| SRR1301346 | SRR1301348 | SRR1301350 | SRR1301352 | SRR1301353 | SRR1301354 |
| SRR1301356 | SRR1301357 | SRR1301359 | SRR1301361 | SRR1301362 | SRR1301363 |
| SRR1301364 | SRR1301365 | SRR1301366 | SRR1301368 | SRR1301370 | SRR1301372 |
| SRR1301373 | SRR1301374 | SRR1301376 | SRR1301378 | SRR1301380 | SRR1301382 |
| SRR1301384 | SRR1301385 | SRR1301387 | SRR1301389 | SRR1301392 | SRR1301394 |
| SRR1301395 | SRR1301396 | SRR1301398 | SRR1301399 | SRR1301400 | SRR1301403 |
| SRR1301406 | SRR1301407 | SRR1301409 | SRR1301411 | SRR1301413 | SRR1301414 |
| SRR1301416 | SRR1301417 | SRR1301418 | SRR1301422 | SRR1301424 | SRR1301425 |
| SRR1301426 | SRR1301430 | SRR1301432 | SRR1301434 | SRR1301436 | SRR1301438 |
| SRR1301440 | SRR1301441 | SRR1301442 | SRR1301444 | SRR1301446 | SRR1301448 |
| SRR1301450 | SRR1301452 | SRR1301454 | SRR1301457 | SRR1301459 | SRR1301460 |
| SRR1301461 | SRR1301463 | SRR1301465 | SRR1301468 | SRR1301470 | SRR1301471 |
| SRR1301475 | SRR1301477 | SRR1301479 | SRR1301481 | SRR1301483 | SRR1301485 |
| SRR1301486 | SRR1301487 | SRR1301489 | SRR1301491 | SRR1301493 | SRR1301495 |
| SRR1301498 | SRR1301500 | SRR1301502 | SRR1301505 | SRR1301507 | SRR1301508 |
| SRR1301509 | SRR1301511 | SRR1301512 | SRR1301513 | SRR1301515 | SRR1301516 |

| | | | | | |
|---|---|---|---|---|---|
| SRR1301517 | SRR1301520 | SRR1301521 | SRR1301523 | SRR1301524 | SRR1301525 |
| SRR1301527 | SRR1301529 | SRR1301531 | SRR1301533 | SRR1301535 | SRR1301537 |
| SRR1301539 | SRR1301541 | SRR1301543 | SRR1301544 | SRR1301545 | SRR1301547 |
| SRR1301549 | SRR1301551 | SRR1301552 | SRR1301555 | SRR1301557 | SRR1301558 |
| SRR1301559 | SRR1301561 | SRR1301563 | SRR1301565 | SRR1301566 | SRR1301567 |
| SRR1301570 | SRR1301571 | SRR1301573 | SRR1301574 | SRR1301575 | SRR1301577 |
| SRR1301578 | SRR1301579 | SRR1301581 | SRR1301583 | SRR1301585 | SRR1301587 |
| SRR1301590 | SRR1301591 | SRR1301593 | SRR1301595 | SRR1301597 | SRR1301599 |
| SRR1301601 | SRR1301603 | SRR1301607 | SRR1301611 | SRR1301613 | SRR1301615 |
| SRR1301617 | SRR1301618 | SRR1301619 | SRR1301621 | SRR1301623 | SRR1301625 |
| SRR1301627 | SRR1301629 | SRR1301631 | SRR1301633 | SRR1301634 | SRR1301635 |
| SRR1301639 | SRR1301641 | SRR1301644 | SRR1301645 | SRR1301647 | SRR1301649 |
| SRR1301651 | SRR1301652 | SRR1301653 | SRR1301655 | SRR1301657 | SRR1301659 |
| SRR1301661 | SRR1301663 | SRR1301664 | SRR1301665 | SRR1301669 | SRR1301673 |
| SRR1301675 | SRR1301677 | SRR1301679 | SRR1301680 | SRR1301681 | SRR1301683 |
| SRR1301685 | SRR1301688 | SRR1301691 | SRR1301692 | SRR1301694 | SRR1301695 |
| SRR1301697 | SRR1301698 | SRR1301700 | SRR1301701 | SRR1301702 | SRR1301704 |
| SRR1301705 | SRR1301706 | SRR1301709 | SRR1301711 | SRR1301712 | SRR1301714 |
| SRR1301717 | SRR1301718 | SRR1301721 | SRR1301722 | SRR1301724 | SRR1301725 |
| SRR1301727 | SRR1301728 | SRR1301729 | SRR1301732 | SRR1301733 | SRR1301736 |
| SRR1301738 | SRR1301740 | SRR1301742 | SRR1301743 | SRR1301744 | SRR1301748 |
| SRR1301750 | SRR1301752 | SRR1301758 | SRR1301759 | SRR1301761 | SRR1301762 |
| SRR1301763 | SRR1301765 | SRR1301767 | SRR1301769 | SRR1301770 | SRR1301771 |
| SRR1301773 | SRR1301775 | SRR1301777 | SRR1301778 | SRR1301782 | SRR1301783 |
| SRR1301786 | SRR1301788 | SRR1301789 | SRR1301790 | SRR1301794 | SRR1301796 |
| SRR1301797 | SRR1301801 | SRR1301803 | SRR1301805 | SRR1301807 | SRR1301809 |
| SRR1301811 | SRR1301812 | SRR1301813 | SRR1301815 | SRR1301817 | SRR1301819 |
| SRR1301821 | SRR1301825 | SRR1301827 | SRR1301828 | SRR1301829 | SRR1301833 |
| SRR1301835 | SRR1301837 | SRR1301840 | SRR1301841 | SRR1301845 | SRR1301848 |
| SRR1301849 | SRR1301851 | SRR1301853 | SRR1301856 | SRR1301858 | SRR1301859 |
| SRR1301860 | SRR1301863 | SRR1301864 | SRR1301867 | SRR1301868 | SRR1301872 |
| SRR1301876 | SRR1301880 | SRR1301883 | SRR1301885 | SRR1301886 | SRR1301887 |
| SRR1301889 | SRR1301891 | SRR1301894 | SRR1301898 | SRR1301902 | SRR1301904 |
| SRR1301906 | SRR1301909 | SRR1301910 | SRR1301914 | SRR1301916 | SRR1301918 |
| SRR1301921 | SRR1301922 | SRR1301926 | SRR1301929 | SRR1301930 | SRR1301934 |
| SRR1301938 | SRR1301940 | SRR1301942 | SRR1301945 | SRR1301947 | SRR1301948 |
| SRR1301949 | SRR1301951 | SRR1301952 | SRR1313245 | SRR1313246 | SRR1313247 |
| SRR1313250 | SRR1313252 | SRR1313254 | SRR1313256 | SRR1515929 | SRR1515931 |
| SRR1515933 | | | | | |
